# Laboratory experiments suggests limited impact of increased nitrogen deposition on snow algae blooms

**DOI:** 10.1101/2024.07.24.604882

**Authors:** Pablo Almela, James J. Elser, J. Joseph Giersch, Scott Hotaling, Victoria Rebbeck, Trinity L. Hamilton

## Abstract

Snow algal blooms decrease snow albedo and increase local melt rates. However, the causes behind the size and frequency of these blooms are still not well understood. One factor that is likely contributing is nutrient availability, specifically nitrogen (N) and phosphorus (P). However, the nutrient requirements of the taxa responsible for these blooms is not known. Here, we assessed the growth of three commercial strains of snow algae under 24 different nutrient treatments that varied in both absolute and relative concentrations of N and P. After 38 days of incubation, we measured total biomass and cell size and estimated their effective albedo reduction surface (EARS). Snow algal strains tended to respond similarly and achieved bloom-like cell densities over a wide range of NP conditions. However, the molar ratio of N:P at which maximum biomass was achieved was between 4 and 7. Our data indicate a high requirement for P for snow algae and suggest that additional N inputs into the ecosystem may not significantly impact the productivity and abundance of snow algae blooms. This highlights P availability as a critical factor influencing the frequency and extent of snow algae blooms and their potential contribution to snow melt through altered albedo.

## Observation

### Significance of studying snow algae

Snow algae are key drivers of biogeochemical cycles in alpine and polar snowfields, as they dominate primary production in these ecosystems (e.g., Lutz et al., 2014; Hamilton and Havig, 2017; Ganey et al., 2017). Under suitable conditions, these microalgae produce colorful snow, with blooms appearing in green, pink, orange, or red as the algae produce photoprotective pigments as adaptive mechanisms for survival and reproduction (Dial et al., 2018). These blooms decrease snow albedo, even when they occur beneath the surface (Almela et al., 2024) and accelerate melt over vast areas (e.g., Engstrom et al., 2023). The rate of snowmelt is crucial as snowpack provides water for one-sixth of the global population (Barnett et al., 2005), with many communities relying on mountain glacier meltwater for agriculture, hydropower, and drinking water (Milner et al., 2017). Therefore, understanding the factors driving these algal blooms is relevant for managing water resources and predicting ecosystem changes.

### Nutrients in the alpine environment

Polar and high-altitude (alpine) mountain regions are typically cold, strongly irradiated, and nutrient-poor environments. In particular, surface ice and snow have lower nutrient concentrations compared to other microhabitats in the same ecosystems (Ren et al., 2019). The input of carbon (C) into the system takes place mostly through CO_2_ fixation, where primary production promotes the accumulation of autochthonous organic C (e.g., Havig and Hamilton, 2019). Nitrogen (N) and phosphorus (P) are two of the most commonly limiting elements for primary production (Elser et al. 2007; including in snow Stibal et al. 2009), and their availability can strongly influence population dynamics, community structure, and ecosystem processes. While community-level primary production is typically constrained by multiple nutrients, an imbalance in a single nutrient can significantly impact freshwater and terrestrial ecosystems (Elser et al. 2007). Limited data from previous studies have indicated an N:P ratio of 11 to 20 (molar, and hereafter) on snowfields where snow algae are abundant (Spijkerman et al., 2012). However, it is unclear what N:P ratio is ideal for snow algae growth, or if interactions between snow algal growth and N or P availability varies among species. Better understanding of the N/P requirements of snow algae will help in assessing how imbalances in N or P availability impact snow algae bloom development.

### Results

We performed laboratory growth experiments to assess the preferred NP ratio for growth of snow algal blooms. The dominant taxa observed in snow algae blooms are represented by three genera: *Chlamydomonas, Chloromonas*, and *Sanguina* (Hotaling et al., 2021). We selected three commercially available representative snow algae strains: *Chlamydomonas augustae* SN134 (Tioga Pass, California), *Chloromonas rosae* UTEX B SNO65 (Litchfield Island, Antarctica) and *Chloromonas typhlos* CCAP11/128 (Sierra Nevada, California) (renamed from *Chlamydomonas nivalis* after Procházkova et al. 2019). There are no commercial strains of *Sanguina* currently available.

We conducted our experiment with factorial design in 24-well plates with each well containing 2 mL of Modified Bold 3N Medium (MB3M). MB3M was prepared following the UTEX Culture Collection of Algae (UT-Austin, USA) recipe, except for the concentrations of N and P. Treatments were allowed to incubate for 38 days at 4.5°C. For comparison, eutrophic lakes generally exhibit total N concentrations ranging from 650 to 1,200 μg L^−1^ and total P concentrations from 30 to 100 μg L^−1^ (Dodds and Whiles, 2010). In contrast, snow and ice on mountain glaciers typically contain a fraction of these levels (Ren et al., 2019). In this experiment, the N and P levels spanned a spectrum from oligotrophic to eutrophic conditions, mirroring the nutrient availability observed in snow fields that support snow algae (e.g., Spijkerman et al., 2012). Complete Materials and Methods are provided in Supplemental Information.

Snow algae strains responded differently to various N:P ratios but generally grew better under N:P ratios between 5.6 and 8.1 in early stages (days 14 and 18). By the end of the experiment (day 38), the highest cell densities, estimated from absorbance (R>0.93, p<0.05), were observed across a wider range of N:P ratios, from 4.2 to 17.5, although the peaks were between 4.2 and 7.2. The differences observed throughout the study may be explained by whether the strains are in their exponential or equilibrium growth phases, a key determinant of their optimal stoichiometry (Klausmeier et al., 2004). Cell densities differed significantly among the three snow algae strains when considering the nutrient treatments with the highest response. *Chloromonas typhlos* reached the highest maximum cell density (NP ratio of 7.20) with ∼150 × 10^5^ cells mL^−1^, which is 3 to 5 times greater than the maximum densities observed in *Chloromonas rosae* (NP ratio of 5.60) (36 × 10^5^ cells mL^−1^) and *Chlamydomonas augustae* (NP ratio of 4.20) (54 × 10^5^ cells mL^−1^) (**Figure 1A**).

**Figure 1.**
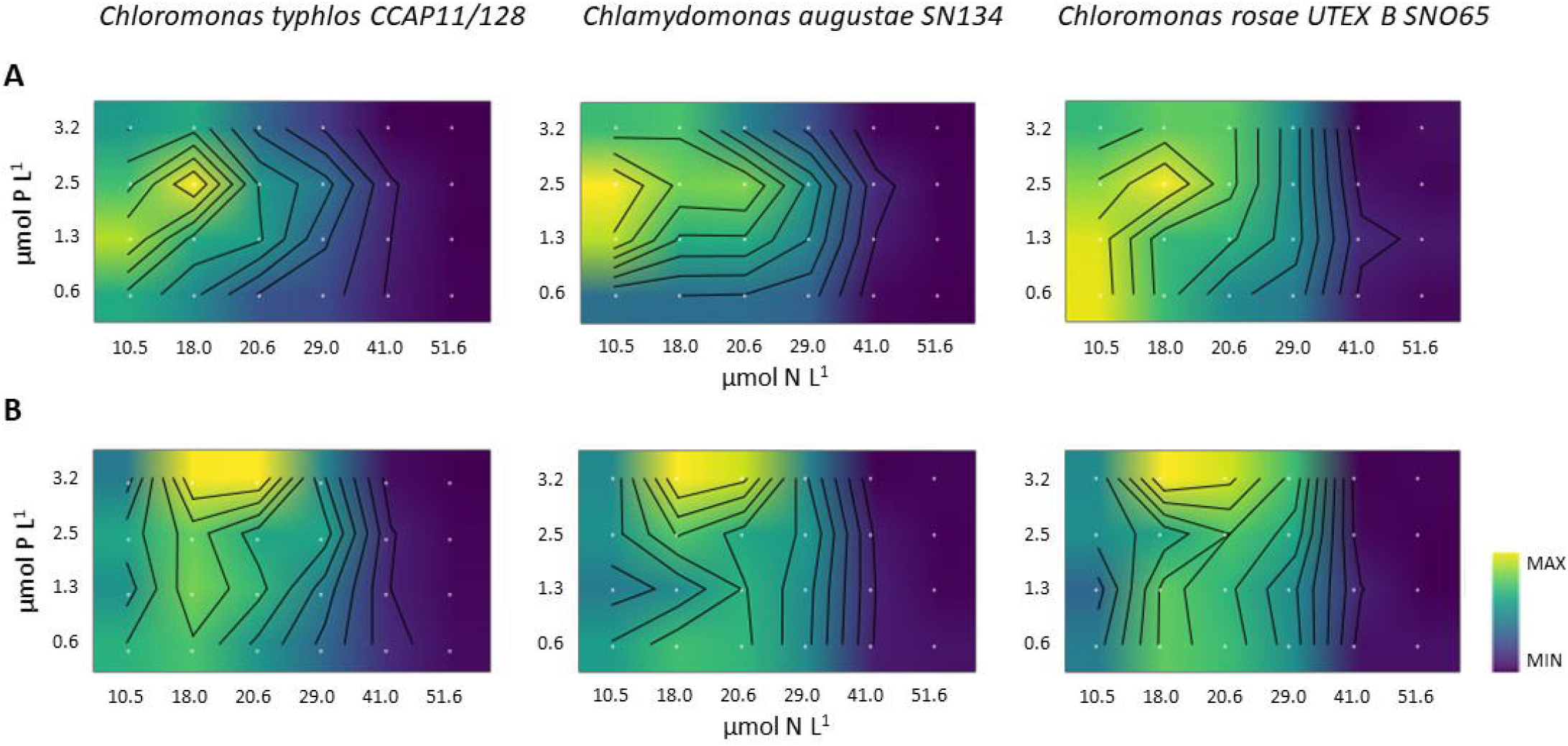
Heat map for (**A**) cell densities and (**B**) chlorophyll concentrations across varying NP treatments. Yellow indicates highest concentrations, while blue indicates lowest concentrations for each strain independently. The x-axis displays nitrogen concentrations, while the y-axis represents phosphorus concentrations (μmol L^−1^).

*Chloromonas typhlos* did not show a response (>10^4^ cells mL^−1^) for two of the 24 nutrient treatments, and *Chlamydomonas augustae* for six, whereas *Chloromonas rosae* showed a positive response for all nutrient treatments. For reference, snow algae blooms typically show concentrations of around 1-20 × 10^5^ cells mL^−1^ (e.g., Müller et al., 1998), although some studies report significantly lower densities (10^3^-10^4^ cells mL^−1^; Lutz et al., 2016). When considering chlorophyll concentrations (**Figure 1B**), the N:P supply ratios resulting in maximum biomass for *Chloromonas typhlos, Chloromonas rosae*, and *Chlamydomonas augustae* were between 5.6 and 6.4. While these findings cannot be directly applied to natural snow conditions, they suggest that snow algae are able to grow under a wide range of nutrient levels, enabling them to proliferate in snow environments ranging from oligotrophic to eutrophic.

N:P ratios also significantly influenced cell area across treatments and species (p<0.0001), with lower density treatments exhibiting cells twice the area of those in higher density treatments (**Figure 2**). Nutrient stress, such as N and P limitation, can cause algae to reduce cell division and accumulate carbohydrates, lipids, or polyphosphate bodies, leading to larger cell sizes (Yan et al., 2021; Grover, 1989). Stress can also induce the synthesis of secondary carotenoids in snow algae (e.g., Leya et al., 2009). For instance, species from the genera *Chlamydomonas* and *Chloromonas* (Chlorophyceae) produce these carotenoids during vegetative and resting stages (e.g., Remias *et al*., 2005), allowing them to maintain photosynthesis under stress as secondary carotenoids stabilize pigment–protein complexes. Although nitrogen limitation is considered a potential trigger for photoprotective carotenoid production in snow algae (Leya et al., 2009), we observed no obvious differences in pigmentation across treatments or algae strains, although we did not quantify carotenoid pigments in this study.

**Figure 2.**
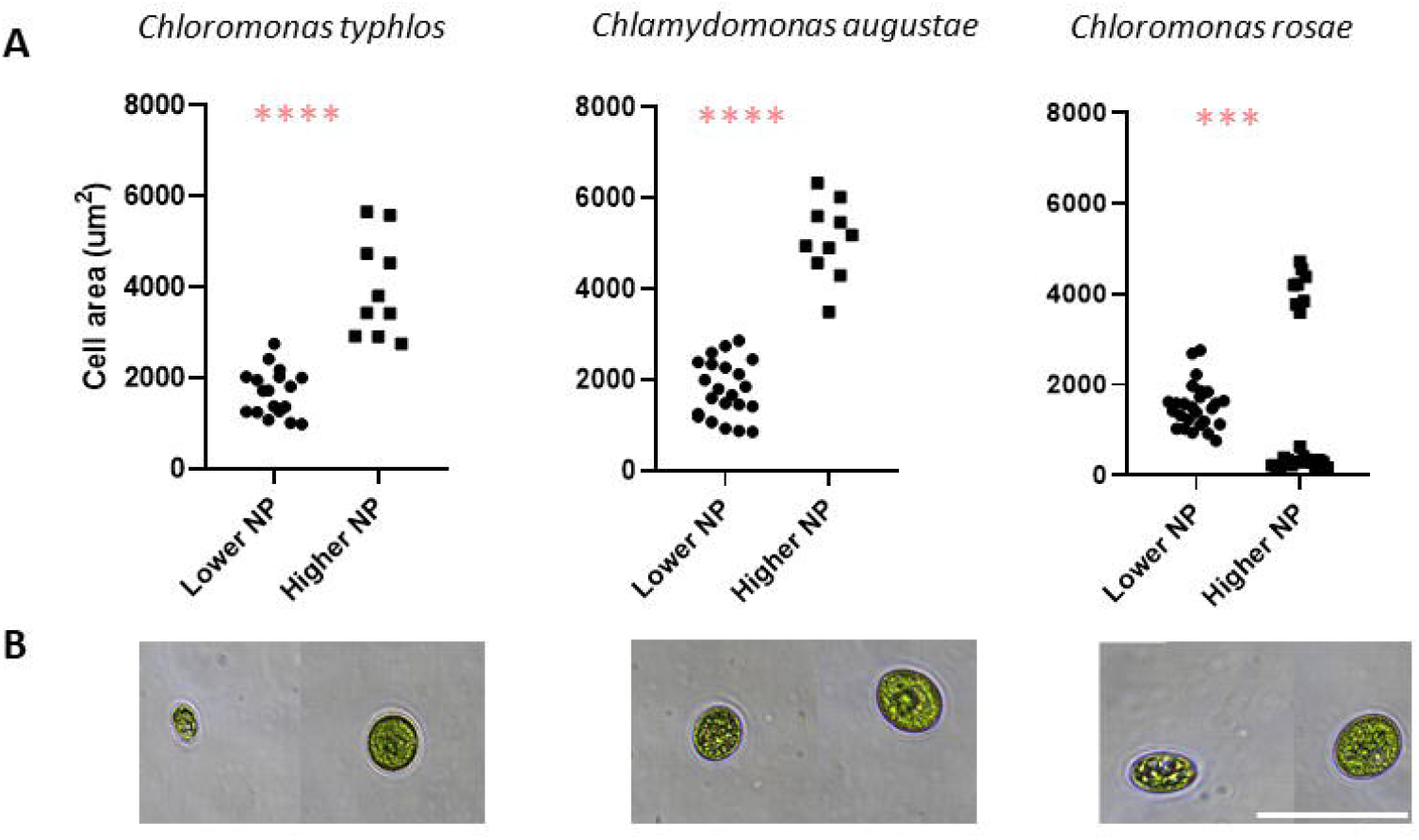
(**A**) Scatter plot illustrating variations in cell area (μm^2^ cell^−1^) of selected nutrient treatments with the highest cell densities (lower N:P = 7.2 and 8.1) compared to those with the lowest cell densities (higher N:P = 16.4 and 31.5) for *Chloromonas typhlos* (*C. nivalis*), *Chlamydomonas augustae* and *Chloromonas rosae*. Significance of comparisons indicated by asterisks after Mann-Whitney test. (**B**) Microscope images demonstrating differences in cell size between the aforementioned nutrient treatments (scale bar = 50 μm).

We found a positive correlation between cell density (cells mL^−1^) and chlorophyll-a concentration for all three snow algae strains (R=0.6, p<0.05), indicating a similar chlorophyll per cell value regardless of nutrient treatment. The biological albedo reduction (BAR) capacity of snow algae is thought to be associated with cell density and pigment content (Hotaling *et al*. 2021). Therefore, we assessed potential differences in cell size on the potential energy absorption capacity and their subsequent impact on albedo reduction for these species. Using cell density and the known cell area, we estimated a metric we refer to as the effective albedo reduction surface (EARS). We evaluated EARS by comparing differences between the most and least favorable nutrient treatments for the three snow algae strains. For *Chloromonas typhlos*, the differences between high-response (high P) and low-response (low P) treatments when considering biomass were substantial, with cell density differing by a factor of 10 and chlorophyll concentration by a factor of 9. However, when accounting for the effective surface area, these differences were substantially reduced, differing by a factor of 4 (**Table S2**). A similar trend was observed for *Chloromonas rosae*. Therefore, the differences in EARS between the most and least favorable nutrient treatments were reduced compared to the differences observed when considering only their biomass (i.e., cell density or chlorophyll concentration). This suggests that EARS could be a valuable indicator for measuring the capacity of snow algae to reduce albedo, as it accounts for both cell density and cell size. While pigments play a crucial role in absorbing sunlight and converting it to heat, a larger surface area may have a similar effect even with lower pigment concentration.

Despite the differences obtained between the two methods for assessing biomass response under the broad range of nutrient conditions (**Figure 1**), our results indicated that snow algae strains tend to grow best at N:P molar ratios lower than 9, which is less than half of the classic Redfield ratio of N:P = 16 (Redfield, 1958). Seasonal snowfields in alpine ecosystems offer a brief growing season that benefit fast-growing biota, which typically exhibit low C:P and N:P ratios due to their higher allocation to P-rich ribosomal RNA (Elser et al., 2003). Therefore, this disproportionate allocation to P suggests that P availability is likely to be a limiting factor for the growth of snow algae across a broad range of conditions. Our results corroborate previous findings showing that P has a pivotal role in shaping microbial dynamics and ecological processes in cold environments (Stibal et al. 2009, 2008). Agriculture and fossil fuel combustion have led to increasing N emissions into the atmosphere (Li et al., 2016), resulting in high N:P ratios upon deposition. Our findings suggest that if N input to snow is not accompanied by an increase in P—an element that is likely more limiting to snow algae in these ecosystems and less bioavailable to the community—it may not significantly influence the productivity and abundance of snow algae. However, extreme wildfire events, which are becoming more frequent in the western U.S., can significantly contribute to nutrient deposition (Campbell et al., 2022) and may increase P bioavailability in alpine ecosystems (Allin et al., 2012). For instance, Spencer et al. (2003) reported a 5 to 60-fold increase in NO_3_^−^, NH_4_^+^, and PO_4_^3−^ concentrations above background levels in streams following a wildfire in Glacier National Park, Montana. Therefore, increased N deposition due to fossil fuel combustion or volatilized fertilizer components may not be a primary driver of more intense snow algae blooms. Instead, the disproportionate requirement for P that our data indicate for snow algae suggests that the rising frequency and severity of wildfires—and their impact on P deposition—could significantly alter nutrient dynamics and ecosystem productivity in high alpine ecosystems. This could promote the frequency and extent of snow algae blooms and increase snow melt through altered albedo, not just from wildfire ash itself but also from snow algae pigments resulting from algal growth in response to ash-deposited phosphorus.

## Author contribution

PA and TH designed the study. PA conducted the experiments and the laboratory analysis. PA wrote the initial manuscript. All authors contributed to interpreting the data and writing the final manuscript.

## Acknowledgements

We would like to thank W. Harcombe and J. Bazurto for generously providing access to their plate readers, which was essential for conducting our experiments.

