## Supplemental Information for "Laboratory experiments suggests limited impact of increased nitrogen deposition on snow algae blooms"

**Material and Methods**

*Snow algae species*

We conducted our experiment using algae strains previously described or isolated from snow. These 3 strains are: *Chlamydomonas augustae* SN134 (isolated from Tioga Pass, California, USA), *Chloromonas rosae* UTEX B SNO65 (isolated from Litchfield Island, Antarctica) and *Chloromonas typhlos* CCAP11/128 (isolated from Sierra Nevada, California, USA). We obtained the cultures from the UTEX Culture Collection of Algae at UT-Austin (US) and the Culture Collection of Algae and Protozoa (UK). Algae were pre-grown in cell culture flasks for several months, using sterile MB3M medium which was replaced every month prior to the experiment, at 4.5°C with a 12:12 h light:dark cycle (30 μmol PAR m⁻² s⁻¹ ≈ 6.5 W m⁻²).

*Experimental design, medium and nutrient treatments*

We conducted a 38-day laboratory factorial experiment in 24 well plates containing 2 mL of medium each. The medium was based on the Modified Bold 3N Medium (MB3M), but with different concentrations of N and P compared to the full medium. The supplied concentrations were 10.5, 18.0, 20.6, 29.0, 41.0 and 51.6 μmol N L^−1^ and 3.2, 2.5, 1.3 and 0.6 μmol P L^−1^, yielding 24 possible combinations (i.e., nutrient treatments) of absolute and relative N:P supply (**Table S1**). For all other nutrients, except GR+ Medium which was not added, we used the concentrations of the full MB3M medium, which reduces the risk of limitation by other nutrients than N and P.

|  | N (µmol L^-1^) | | | | |
| --- | --- | --- | --- | --- | --- |
| P (µmol L^-1^) | **10.5** | **18.0** | **20.6** | **29.0** | **41.0** |
| **3.2** | 3.3 | 5.6 | 6.4 | 9.1 | 12.8 |
| **2.5** | 4.2 | 7.2 | 8.2 | 11.6 | 16.4 |
| **1.3** | 8.1 | 13.8 | 15.8 | 22.3 | 31.5 |
| **0.6** | 17.5 | 30.0 | 34.3 | 48.3 | 68.3 |

**Table S1.** Experimental design with N and P supply (μmol L^−1^, bold letters) and molar nutrient ratios for all 24 combinations of N and P supply.

Nutrient treatments and algae were added at the beginning of the experiment and afterward no addition took place. To ensure uniformity in nutrient levels across the various species and treatments, an equal volume of algae from the original culture (50 μL) was inoculated to each treatment at the beginning of the experiment. Our main goal was to examine how each different species responds to the treatments rather than to compare among them, making the initial use of different biomass not crucial. However, all strains had a cell density >10^4^ cells/mL at the time of inoculation. The 24-well plates containing the algae were incubated at 4.5°C under a 12:12 h light-dark cycle with an intensity of 30 μmol PAR m⁻² s⁻¹ (≈ 6.5 W m⁻²).

*Snow algae biomass: cell densities, cell areas and chlorophyll-a concentrations*

Growth of cultures was measured every 2 days by optical density (OD) at 750 nm (Geller et al., 2018) using a Multiskan SkyHigh Microplate Spectrophotometer (ThermoFisher, USA). Before each measurement, the 24 well plates were shaken manually until the cultures were homogenized. OD readings were taken without the lid, and converted to algal cell concentrations using standard curves plotted from the OD of known cell concentrations for each algal species. At the end of the experiment, a 200 μL aliquot was taken of selected samples for cell counts and to measure cell area. An 800 mL aliquot was taken from each nutrient treatment, and the chlorophyll-a concentration in the cultures was measured as a proxy for algal biomass. The samples were filtered onto 0.7 μm pore size 25-mm GF/F filters, and pigments were extracted with methanol to determine chlorophyll-a concentrations by measuring fluorescent intensity of the extracts, using 24-well plates and a plate reader. The method used is further detailed by Warren (2008).

*Effective Albedo Reduction Surface (EARS) of the snow algae strains*

Using cell density and the known cell area for the selected nutrient treatments, we estimated the Effective Albedo Reduction Surface (EARS) (**Table S2**). We assessed EARS's potential contribution by comparing the differences between the most favorable and least favorable nutrient treatments for the three snow algae strains, and then comparing these differences to those in cell density and chlorophyll-a concentration under the same nutrient conditions.

*Statistical analysis*

We used Pearson correlations to assess relationships between cell densities and chlorophyll-a concentrations. We conducted a Mann Whitney test for evaluating the differences in cell area between the selected nutrient treatments.

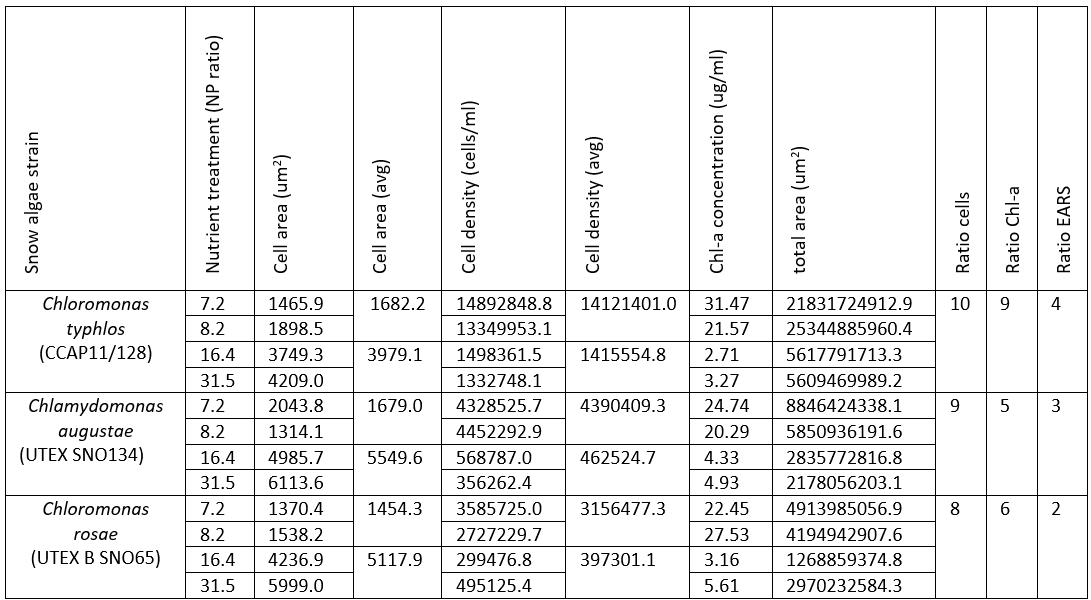

**Table S2.** Summary of cell area, cell density, total area (EARS), total chlorophyll-a concentrations, and ratios of cell densities, chlorophyll-a concentrations and EARS across the selected nutrient treatments for the three snow algae strains.
